# Dieback of stringybark eucalypt forests in the Mount Lofty Ranges

**DOI:** 10.1101/2022.11.03.515134

**Authors:** Gregory R Guerin, Gunnar Keppel, Stefan Peters, Amelia Hurren

## Abstract

Canopy dieback and concerning rates of tree mortality have been noted in iconic forests of the Mount Lofty Ranges (MLR), South Australia, dominated by the stringybark eucalypt species *Eucalyptus baxteri* (Brown Stringybark) and *E. obliqua* (Messmate Stringybark). The extent and causes of stringybark forest decline are not yet fully understood, prohibiting evidence-based management strategies. Here, we explore the distribution of MLR populations of the two species and their position in climate space relative to eastern populations. We also conducted field assessments to investigate stand health and dieback aetiology, and analysed existing tree monitoring data. Stringybarks in the MLR are disjunct from eastern populations and occupy a more summer-arid niche. The species are also susceptible to summer water stress and *Phytophthora*. Periods of drought during 2006–2009 and 2018–2019 may have contributed to observed dieback. However, field assessments suggest a complex landscape syndrome that includes borer infestations and fire impacts among other factors, rather than solely hydraulic failure. Messmate Stringybark has suffered widespread but patchy stand collapse. There is no obvious common pattern of collapsed sites with respect to topography or local water availability (e.g., swamps and ridges equally affected), although northern range-edge sites are heavily affected. Brown Stringybark is less affected but has notable collapse sites. We hope these studies establish a springboard for future investigations and more widespread sampling of MLR stringybark forests. Further investigations should include regional surveys of stringybark sites to record spatial and temporal patterns of tree mortality combined with multi- or hyperspectral analysis of remotely sensed imagery and visual inspection of dieback from very high resolution aerial images and ground-truthing. Our findings confirm the susceptibility of stringybark forests in the MLR to ecosystem collapse and highlight the urgent need to understand the causes and aetiology of the observed dieback.

## Introduction

Dieback events (DEH 2004) associated with drought and extreme heat conditions are being observed in forests globally (Schuldt et al. 2020). In Australia, climate change is implicated as a major driver of tree mortality, although the impacts of drought on forests and the role of other stressors are still poorly understood in many cases (White 2015; Sarnow 2021). Episodes of dieback involving excessive canopy loss or premature death in eucalypts have been observed in Australia since the 1800s (Jurskis 2005). Tree decline accelerated in the late Twentieth Century associated with insect outbreaks and environmental changes (Jurskis 2005; Croft 2017). More recently, mass-mortality events have been observed globally and in Australia, linked to heat waves and drought (Vanassche 2011; De Kauwe et al. 2020; Pappas et al. 2022).

Climate change is implicated as a major driver of dieback linked to hydraulic failure or pest outbreaks (Hoffmann et al. 2019; Nolan et al. 2021; Pappas et al. 2022). However, the aetiology of dieback and the impacts of altered water regimes on Australian forests remain poorly known (Ward 2005; Croft 2017; De Kauwe et al. 2020). Given the importance of forests to the ecosphere, there is an urgent need to assess the extent and aetiology of tree decline (Hammond et al. 2019). For example, under what conditions does hydraulic impairment occur (Hoffmann et al. 2011; De Kauwe et al. 2020; Nolan et al. 2021) and how do soil depth, functional traits and tree density modulate the response (Blackman et al. 2019)?

Forests of two stringybark eucalypts examined here, *Eucalyptus baxteri* (Benth.) Maiden & Blakely ex J.M.Black (Brown Stringybark) and *E. obliqua* L’Hér (Messmate Stringybark), in the Mount Lofty Ranges (MLR) of South Australia are putatively threatened by increasing frequencies of drought and heat wave conditions linked to climate change. Dieback events have left once dense forest canopy in poor condition and resulted in high tree mortality rates, as reported anecdotally. In extreme cases, entire stands of trees have died while in other patches, trees remain healthy.

### Aetiology – pests, diseases and drought

When exposed to soil moisture deficits, trees limit water loss via transpiration by closing the stomatal pores on leaf surfaces, thereby increasing stomatal resistance (Vanassche 2011). Water loss may continue due to imperfect cuticular resistance. Therefore, water-stressed trees may abscise leaves or branches. Excessive vapour pressure deficits can lead to xylem embolism, reducing hydraulic conductance and ultimately causing hydraulic failure (Hammond 2019; Nolan et al. 2021; Pritzkow et al. 2021). Plant functional traits modulate and reflect these responses to aridity in trees (Ottaviani & Keppel 2018). For example, smaller leaves with higher mass per area are adaptations to more arid environments, while foliar stable carbon isotope abundance (δ^13^C) is related to water use efficiency (Steane et al. 2017).

While climate is implicated in the decline of stringybark forests in the region, it is by no means the only candidate driver of observed dieback. A range of pests and diseases are known to cause episodic dieback (Department for Environment and Heritage 2005; Hanold *et al*. 2006; Hoffmann et al. 2019). *Eucalyptus baxteri* and *E. obliqua* are among the most susceptible eucalypts in South Australia to the notorious pathogen *Phytophthora cinnamomi* (‘PC’), with systematic inoculation leading to nearly 100% mortality rates in *E. baxteri* (Hook 2011; Kueh et al. 2012). Indeed, there are numerous suspected and confirmed PC infestations in the study region in and around areas dominated by stringybark forest (Fig. S9), and PC has been confirmed via laboratory tests in samples from at least one known stringybark dieback site in the region.

Lerps and borers are among insects involved in tree dieback syndromes. An outbreak of bark beetles caused extensive pine mortality in the Rock Mountains of the USA between 1996 and 2008 (Pappas et al. 2022). Seaton (2012) found high levels of borers in sick trees of ‘collapsed’ (i.e., high mortality and >50% canopy loss) eucalypt stands in Western Australia when compared to a background of low levels of borers in sick trees in mainly healthy stands. This suggests some kind of localised feedback loop that promotes borer activity and weakens stands (White 2015). *Phoracantha* longicorn beetles (especially *P. semipunctata* and *P. recurva*), are typically involved in borer infestations in eucalypts. Although these species usually attack trees that are already stressed, they are more efficient at finding stressed trees where whole stands are stressed (Seaton 2012). Drought stress is significant to infestation potential, as dry wood attracts borers, and water-stressed trees may be also less able to produce kino in defence, although assumption needs further testing (White 2015). The causal direction can be ambiguous, that is, whether borers cause tree collapse or are a symptom of existing stress. In either case, the feeding galleries of borers can cause hydraulic failure and tree death when densities are high.

Birds are a natural predator of longicorn beetles (Seaton 2012), yet many woodland birds are in decline across stringybark and other forests in the study region (Prowse et al. 2021). It is plausible that declines in insectivorous bird populations exacerbated by habitat loss and increasing fire frequency (Prowse et al. 2017) and trees weakened by warming or drought stress may be contributing to outbreaks of beetles at patch or landscape scale (Pappas et al. 2022). Adult and larval beetles are a component of the diet of Varied Sittellas, for example (Noske 1985). There are potential feedback loops, in which tree decline impacts insectivorous birds, while borers benefit from reduced predation and trees weakened by fragmentation, disease, fire and drought.

### Aims

Here, we report an investigation into the extent of, and factors contributing to, dieback in Brown and Messmate Stringybark forests in the Mount Lofty Ranges in South Australia. The investigation was supported by a Steering Group with government and NGO representatives to enable policy translation and potential trials of management interventions. Further survey work is still required (through both remote sensing and ground surveys) to fully characterise stringybark dieback in the region.

We addressed the following questions:

1. How does the climatic niche of MLR stringybarks compare to their broader distribution?
2. What is the extent and severity of canopy dieback in MLR stringybarks?
3. What is the aetiology of the dieback? Is there evidence of exacerbating factors such as disease, competition from weeds, borers or lerp infestation?

To address these aims we collated spatio-temporal, environmental and ground survey data, performed field reconnaissance and tree assessment visits to a series of stringybark forest sites and performed desktop analysis including comparison of climatic niches and evaluation of climatic trends for the region.

## Methods

### Stringybarks of the Mount Lofty Ranges

Forests of the stringybark eucalypts *Eucalyptus baxteri* (Brown Stringybark) and *E. obliqua* (Messmate Stringybark) in the Mount Lofty Ranges, South Australia, display varying levels of dieback, observed anecdotally and during local monitoring (e.g., see Guerin et al. 2014; Guerin 2017; Sparrow et al. 2020). Although hydraulic failure during drought periods is suspected in some instances, the extent, timing and cause of these dieback events have not previously been investigated in any detail. MLR stringybark forests occur at the western edge of the species ranges, most likely in genetic isolation from more extensive eastern populations (Biffin et al. 2020). Brown and Messmate Stringybark form extensive, characteristic forests which predominate in high rainfall, elevated landscapes (Fig. 1). These forests support pollination services to adjacent apple and pear orchards via floral resources for bees (Spronk 2017) and are associated with diverse species assemblages in a biodiversity hotspot (Guerin et al. 2016). A recent assessment of the ecologically similar Red Stringybark (*E. macrorhyncha* subsp. *macrorhyncha*) forest in the northern MLR reported serious tree decline related to the interaction of drought with topography (Sarnow 2021).

**Figure 1.**
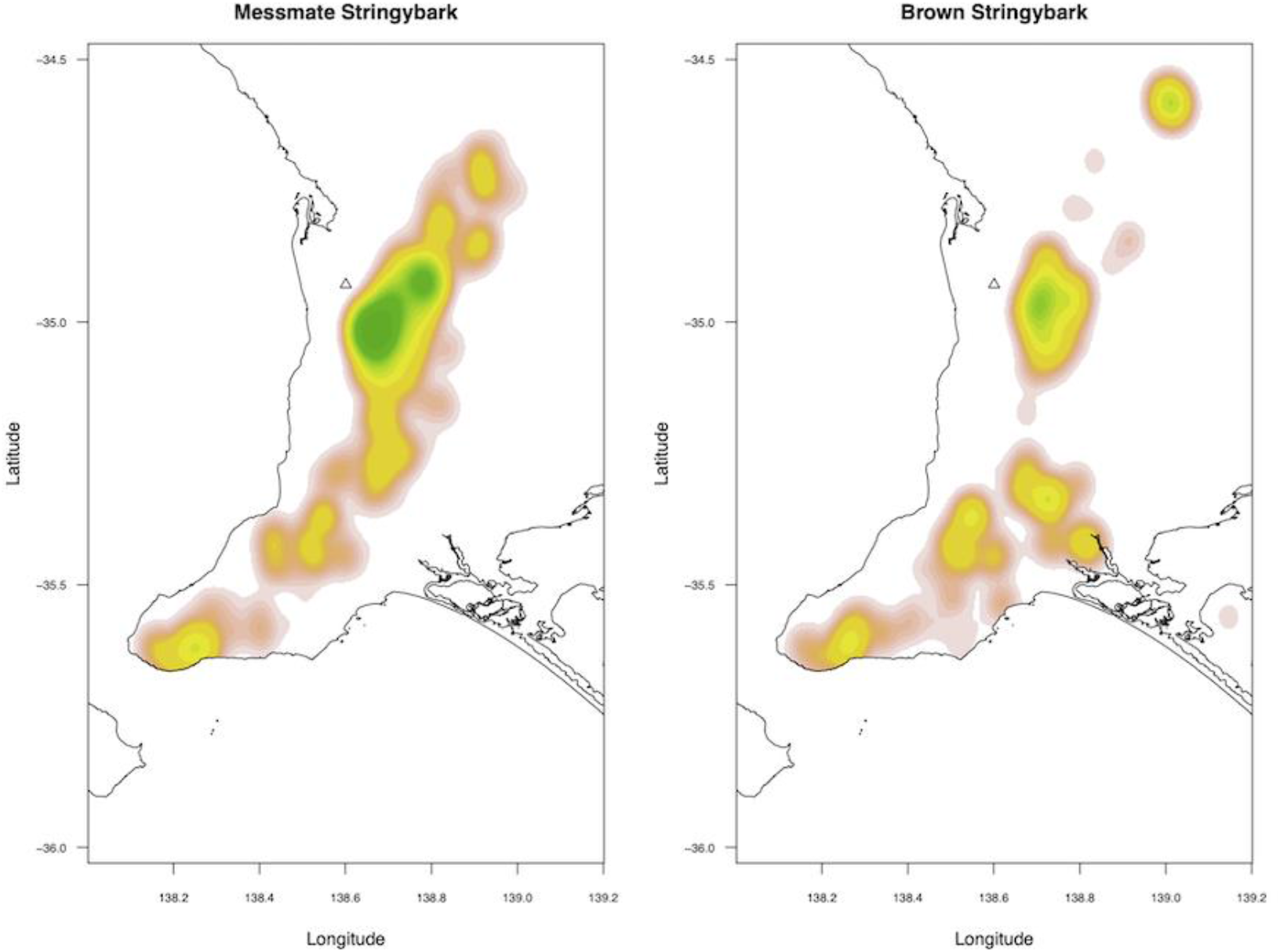
Distribution of Messmate (*Eucalyptus obliqua*; left panel) and Brown (*Eucalyptus baxteri*; right panel) Stringybark in the study region of Mount Lofty Ranges (MLR) in South Australia. Spatial kernel density based on composite presence data. Green fill represents higher spatial densities and highlights the central Adelaide Hills.

### Climatic niche space

Previously recorded presence locations for each stringybark species across their full Australian range were obtained from the Atlas of Living Australia (https://spatial.ala.org.au, accessed 29/3/2021), and subsequently filtered to specimen and observation/survey records with a spatial uncertainty of no more than 1 km. The values of a set of 25 climatic variables at each presence location were extracted from a stack of high resolution (9-second) climate surfaces (Harwood *et al*. 2016). The realised climatic niches of the species were visualised using bivariate scatterplots and by comparing MLR stringybark populations (as well as the adjoining Kangaroo Island) to eastern Australia populations.

### Climatic trends and extremes

Monthly rainfall and mean maximum temperature data were obtained for a selection of weather stations across the geographic range of MLR stringybark and representing areas occupied by the species (http://www.bom.gov.au/climate/data, accessed 16/7/2021). Sites included wetter and drier sites, subject to availability of data throughout the last 20 years as a minimum. We plotted annual rainfall amounts and temperature means as well as monthly values for January and February for each year to assess warming and drying trends and to detect climate pinch-points where low annual/summer rainfall coincided with high maximum temperatures. Rainfall and temperature data were supplemented with drought index data for the period 1950–2022 from SPEIbase, a global database for the Standardised Precipitation Evapotranspiration Index (SPEI) as a drought indicator (Beguería et al. 2014).

### Field reconnaissance and tree assessment visits

We visited a selection of stringybark sites in the region to assess the extent and general condition of stringybark stands, signs of stress, dieback and disturbance, and feasibility of access for monitoring (Table 1). Site visits were selected to cover the geographic range of the species in the study area and to cover high and low rainfall limits of that distribution as well as to include a range of patch conditions, from high quality conservation parks to disturbed local reserves. Site visits were conducted between 2/6/2021 and 8/9/2021.

**Table 1.** Reconnaissance and tree assessment sites visited June–September 2021. Dieback severity codes were defined as follows: *Low* – trees generally healthy with minimal/sporadic canopy loss and stem mortality; *Medium* – trees generally unhealthy with canopy loss/epicormic sprouting or with patchy stem mortality or signs of borer infestation; *High* – 50-80% stem mortality; *Severe* – 80+% stem mortality or with completely collapsed/dead patches.

| Location | Dominant trees | Dieback severity | Dieback notes |
| --- | --- | --- | --- |
| Cleland CP | *E. obliqua* +/- *E. viminalis* +/- *Exocarpos cupressiformis* | Medium | Patchy dead trees. Low to severe borer damage. |
| Mount Pleasant Summit | *E. obliqua* | High | Approximately 50% mortality with borers. |
| Bradwood Soccer | *E. baxteri*/<br>*E. obliqua* | Medium | Some dead trees but many trees infected with borers in early stages. |
| Hahndorf Soccer | *E. obliqua* | Severe | Approximately 80% mortality with borers. Confirmed PC site. |
| Stirling Cemetery | *E. obliqua* | Low | Healthy stands with minimal canopy loss. |
| Lobethal bushland | *E. obliqua* | High | Burnt out in Cudlee Creek fire but historical imagery suggests high mortality. |
| Kaiserstuhl CP | *E. baxteri* | High | Trees in poor condition with dead patches, especially near north-western boundary. |
| Marble Hill–Cherryville | *E. obliqua* | High | Extensive tracts of forest with very high mortality rate. |
| Scott Creek CP | *E. viminalis* +/- *E. obliqua* | Unknown/<br>burnt | Burnt out in the Cherry Gardens fire – anecdotal reports of dieback. |
| Horsnell Gully CP | *E. baxteri*/*E. obliqua* | Medium | Trees unhealthy with extensive epicormic sprouting and canopy loss. Mortality in young stems. |
| Aldgate Tennis | *E. obliqua* | High | Patchy mortality associated with borers. |
| Scott Creek Oval | *E. obliqua* | High | Stringybark dead on lower slope only. Wet, water-logged sites with tall trees. |
| Aldgate Town | *E. obliqua* | High | Patches of stringybarks with high mortality rates. |
| Candlebark Reserve | *E. obliqua*-*E. dalrympeana* | Low | Healthy trees. |
| Vincent Reserve, Bridgewater | *E. obliqua* | Low | Healthy trees with signs of minor borer activity. |
| Carey Gully (White’s Scrub) | *E. obliqua* | Severe | A number of collapsed patches on hillslope and along creek in wet gully – high borer activity. |
| Deep Creek CP | *E. baxteri*/*E. obliqua* | Low | Trees healthy. |
| Eric Bonython CP | *E. baxteri*/<br>*E. obliqua* | Low | Trees healthy, some signs of minor borer activity. |

At each site, we observed tree function and health to develop an understanding of the aetiology of dieback. We describe sites with >50% canopy loss/stem mortality as ‘collapsed’. Tree health inspections recorded presence of canopy dieback, dead trees, and signs and symptoms of stress or damage including leaf burn, basal and epicormic sprouting, mistletoe infestation, significant bark cracking and kino exudation, cankers (fungal infection), canopy thinning, insect attack (e.g., evidence of damage from lerps, skeletonisers, borers and chewers) and pathogens (e.g., chlorosis of leaves, signs in the vegetation of *Phytophthora cinnamomi*).

### Analysis of existing tree monitoring data

We obtained tree data from Bushland Condition Monitoring sites (BCM; Croft *et al*. 2005). BCM includes a tree health module and has been used to survey hundreds of sites in the region and across South Australia. It provides quantitative measures at individual tree level that can potentially be remeasured across a network of sites with existing baselines. Tree data with measurements of alive/dead status, size and canopy extent were obtained from 150 monitoring sites where either of the stringybark species had been recorded. Revisited site data for trees included 82 surveys at 34 sites. The BCM tree method involves a point-centred configuration in which the ten nearest trees to the permanent marker stake of an associated vegetation monitoring plot are scored. Measures include GBH (i.e., girth at breast height) of only the largest stem (where tree is multi-stemmed) and health and habitat scores relating to mistletoe infestation, lerp damage and canopy dieback. While in BCM, these measures are converted to categorical scores, here we considered only the raw data.

Coarse patterns and trends were explored across years by including all BCM tree data, noting that apparent temporal trends could be confounded by the spatial variation in which sites were surveyed in a given year. Trends were explored in more detail for the 34 sites at which the 10 nearest trees to the ‘centre-point’ had been re-measured at least once. For these revisited plots, we plotted baseline–latest visit increments in canopy dieback %, stem mortality and stem basal area (cross-sectional area of living tree trunk per unit area), which accounts for tree growth and mortality.

Basal area from these data are only based on the primary/largest stem of multi-stemmed trees (which are common due to historical logging and fires) as only the largest stem per tree is measured in the standard BCM method. While this metric does not represent true basal area, it still gives some indication of tree growth across a sample of stems. Additionally, trees are not always tagged and can therefore only be identified upon revisit by careful comparison of baseline and revisit distance and bearing from the centre-point. Different trees could then be included in repeat surveys, so that the proportion of living stems can appear to increase when in fact baseline-sampled trees have died and fallen between visits.

### Conceptual model

We developed a conceptual model of MLR stringybark forest dieback aetiology based on literature, climatic niche and event pattens and field observations. The aim was to map complex putative landscape interactions and identify pathways in need of further research.

## RESULTS

### Climatic niche space

MLR populations of both stringybark species represent disjunct, western extremes of the wider Australian distribution of the species (Fig. 2). For a number of climatic gradients, MLR stringybark sit within the climatic niche of the eastern populations, albeit towards the warmer/drier end of that niche (Fig. 3). For example, MLR stringybarks do not occur in habitats with lower mean annual rainfall as compared to the eastern populations. Significantly, however, MLR populations of both species are outliers in terms of summer aridity and its interaction with water deficit. Kangaroo Island (adjacent to MLR) populations of *E. obliqua* and *E. baxteri* are even more outlying in terms of summer aridity, although less exposed to temperature extremes.

**Figure 2.**
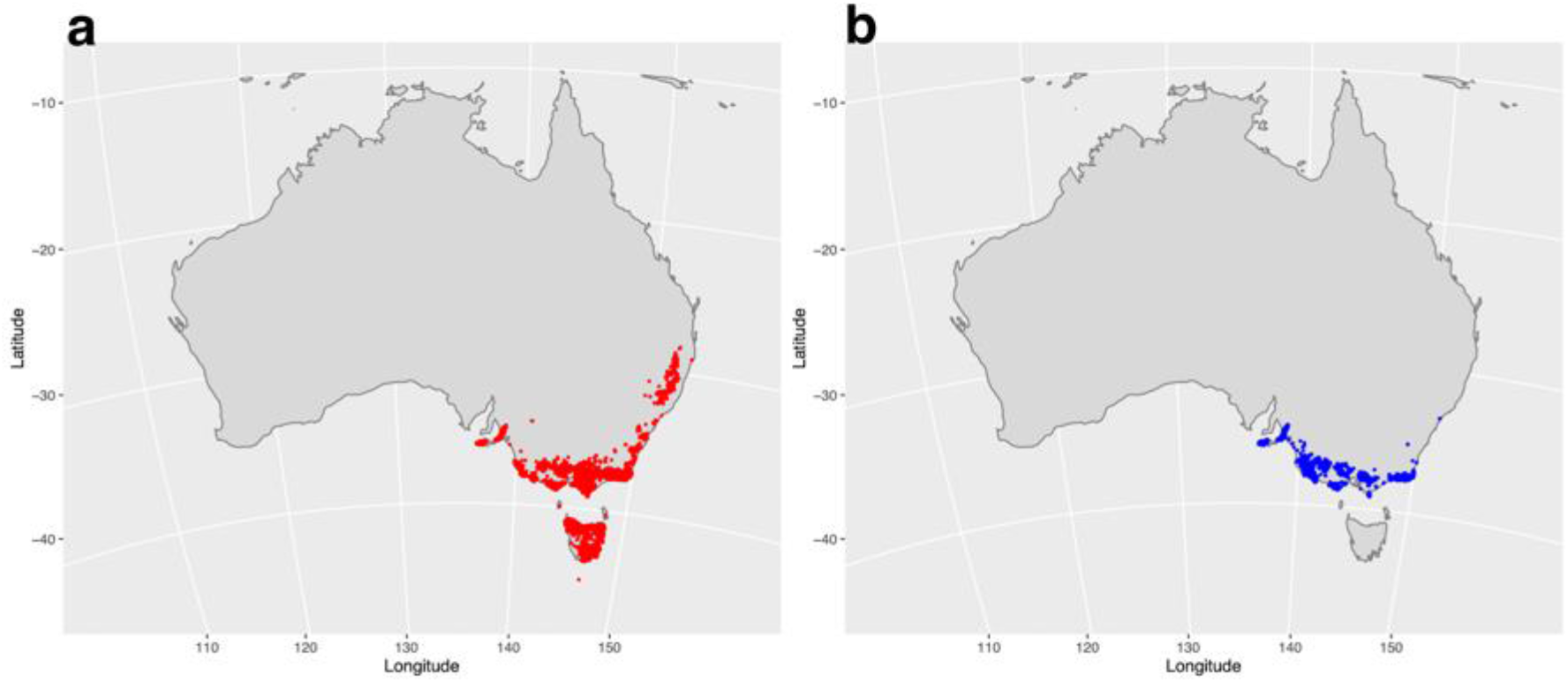
The Australia-wide distribution of a) Messmate Stringybark (*E*. obliqua) and; b) Brown Stringybark (*E. baxteri*). MLR populations (along with adjacent Kangaroo Island populations) are disjunct and the western extreme of the species distributions.

**Figure 3.**
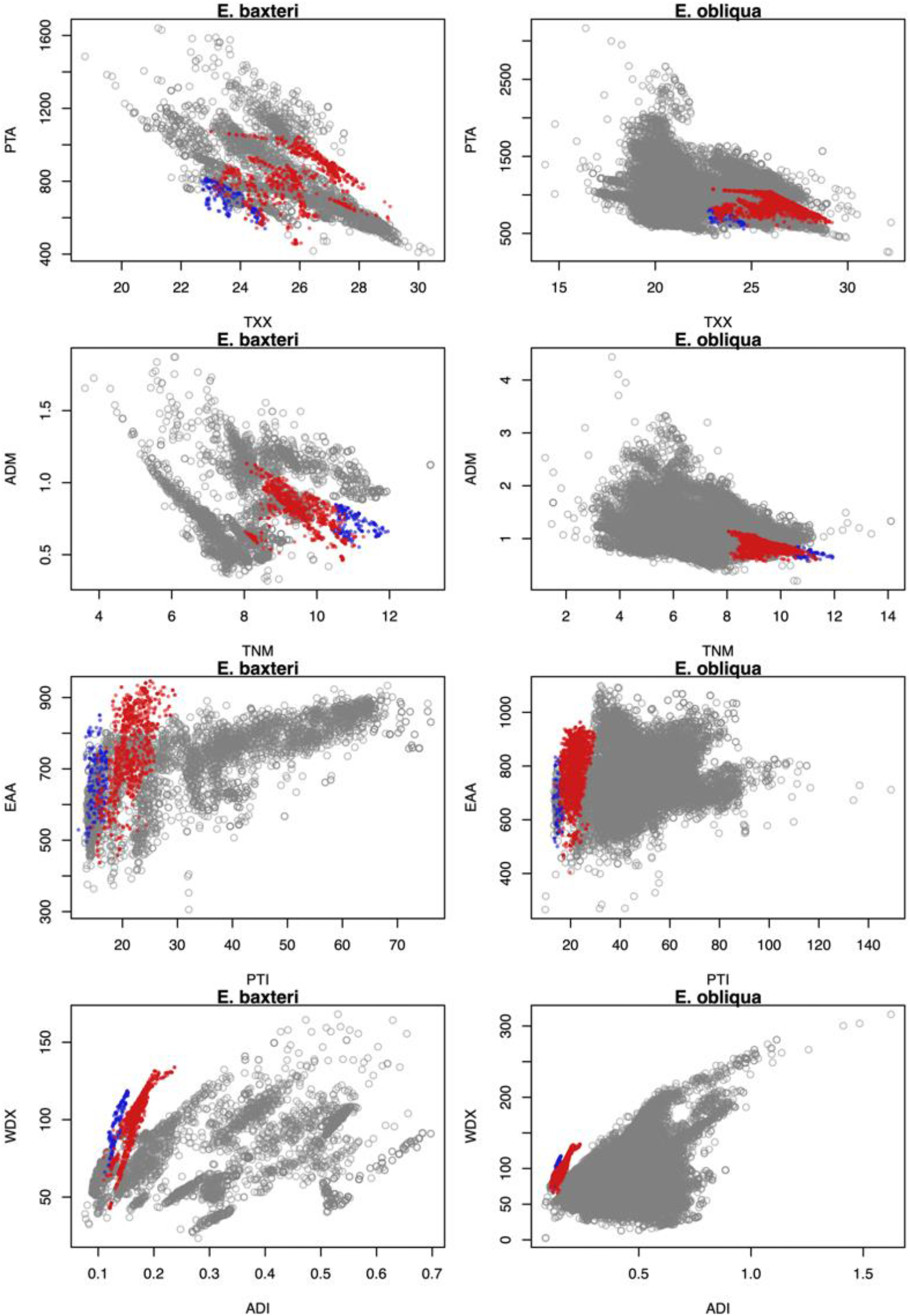
Scatterplots showing the realised climatic niche of MLR (red) and Kangaroo Island (blue) stringybark populations in the context of the species Australia-wide distribution. Left-hand panels = *E. baxteri*; right-hand panels = *E. obliqua*. Axis codes: PTA = Annual precipitation (mm); TXX = Maximum temperature - monthly maximum (°C); ADM = Mean annual aridity index (higher in more mesic environments); TNM = Minimum temperature – Annual mean (°C); EAA = Annual total actual evapotranspiration (mm); PTI = Minimum monthly precipitation (mm); WDX = Minimum monthly atmospheric water deficit (mm); ADI = Minimum monthly aridity index (higher in more mesic environments).

### Climatic trends and extremes

All weather stations in the region followed the same inter-annual patterns, despite sites differing in mean rainfall and temperature (Fig. 4). A significant warming trend of 1.2 degrees has occurred in mean annual maximum temperatures since 1995, with linear models explaining as much as 48% of deviance at Parawa. No significant long-term linear trends were detected in rainfall. Mean maximum temperatures increased for January but decreased for February over the same period, although neither trend was statistically significant due to high interannual variance. The SPEI anomaly has trended negative since 1950 (Fig. S10), signalling more frequent periods of aridity.

**Figure 4.**
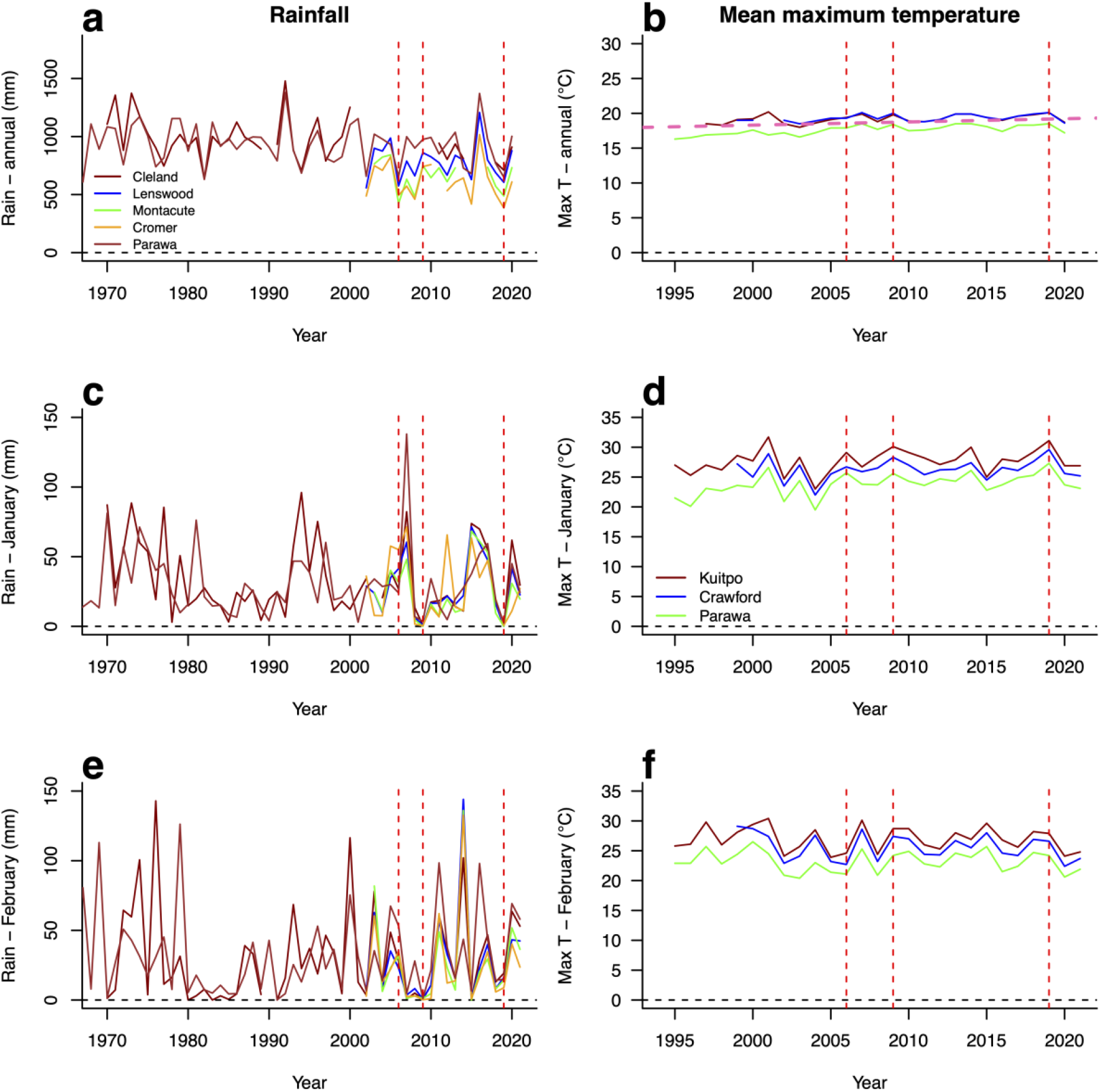
Rainfall and mean maximum temperature data from weather stations in MLR stringybark regions: Cleland, Lenswood, Montacute, Cromer, Parawa, Sevenhill. Dashed red vertical lines highlight drought years which coincided with high summer temperatures (2006, 2009 and 2019).

Since 1970, there have been several significant climatic events, notably drought years with low annual rainfall in 2006, 2015 and 2019 (Fig. 4). The millennium drought involved a dry year in 2006 coinciding with a significant peak in January temperature maxima, followed by dry summers, peaking in January of 2009, during which there was effectively zero rainfall. The dry year of 2015 was perhaps less severe because there was some summer rainfall and the mean of January maxima was not extreme. The year 2019 was a significant event during which low annual rainfall coincided with extremely low summer rainfall and a high peak in January temperature maxima. Both events, 2006–2009 (drought followed by dry summers) and 2019 (drought with dry summer and high temperatures) were likely to have been ecophysiologically significant. SPEI data also show drought periods in 2006–2009 and 2018– 2019, exceeding aridity levels seen in the 1960s and 1980s.

### Field reconnaissance and tree assessment visits

Dead and dying stringybarks were observed across the region, with the exception of a few stands (Table 1; Fig. 5). Tree mortality at sites affected by dieback ranged from a few dead trees scattered among living ones, up to 50–80% mortality at a number of collapsed sites. Indeed, dieback observed at nine of the 18 reported sites was scored as *high* or *severe* while only five were scored as *low*. The timing of observed tree mortality ranges from ‘old’ (age unknown but bark no longer persisting on tree remnants) to very recent deaths, where bark and small twigs are still attached, for example. Additionally, trees were frequently observed to be in the process of dying even where existing stand mortality was relatively low. Of the sites visited, significant mortality was observed at Mount Pleasant (summit), Tanunda (Kaiserstuhl), Hahndorf, Aldgate, Carey Gully and Montacute. Collapsed stands were less evident on the Fleurieu Peninsula in the southern part of the study region, though occasional patches of stringybark had some dead trees.

**Figure 5.**
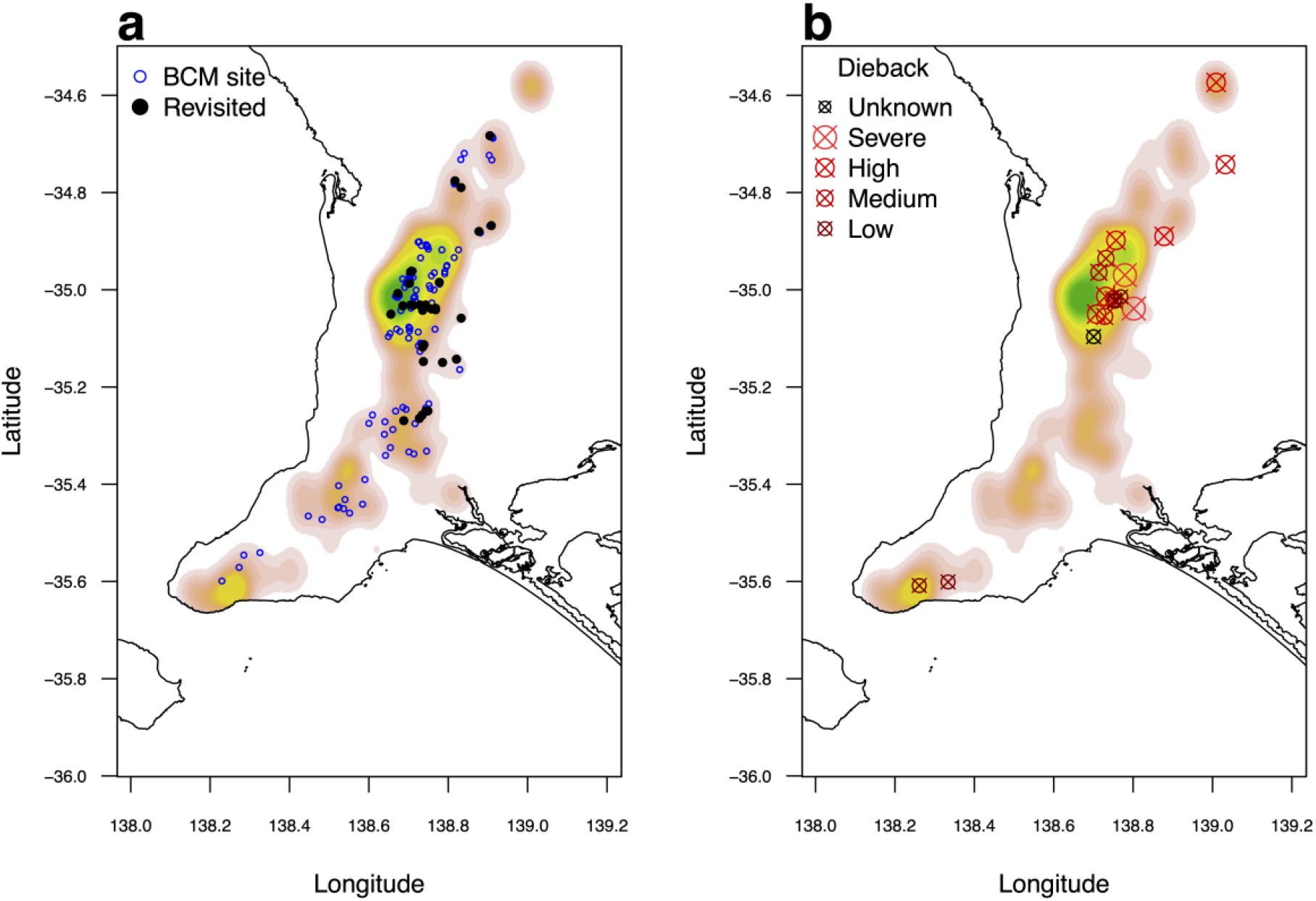
a) Location of Bushland Condition Monitoring (BCM) sites with tree health module data in stringybark forest in the southern MLR (revisited sites highlighted); b) field assessment sites, coded by severity of observed dieback (see Table 1 for codes). Sites are plotted over spatial kernel density map for both stringybark species combined.

Tree mortality was consistently associated with heavy infestations of eucalypt boring longicorn beetles (*Phoracantha spp*.). High densities of oviposition sites, larval galleries, exit holes and frass were almost uniformly found on recently dead and dying trees (Fig. 6). Moreover, unlike attacks from leaf herbivory and recovery post drought and fire, trees infested with wood borers are killed outright and were not observed to resprout from epicormic buds nor basally.

**Figure 6.**
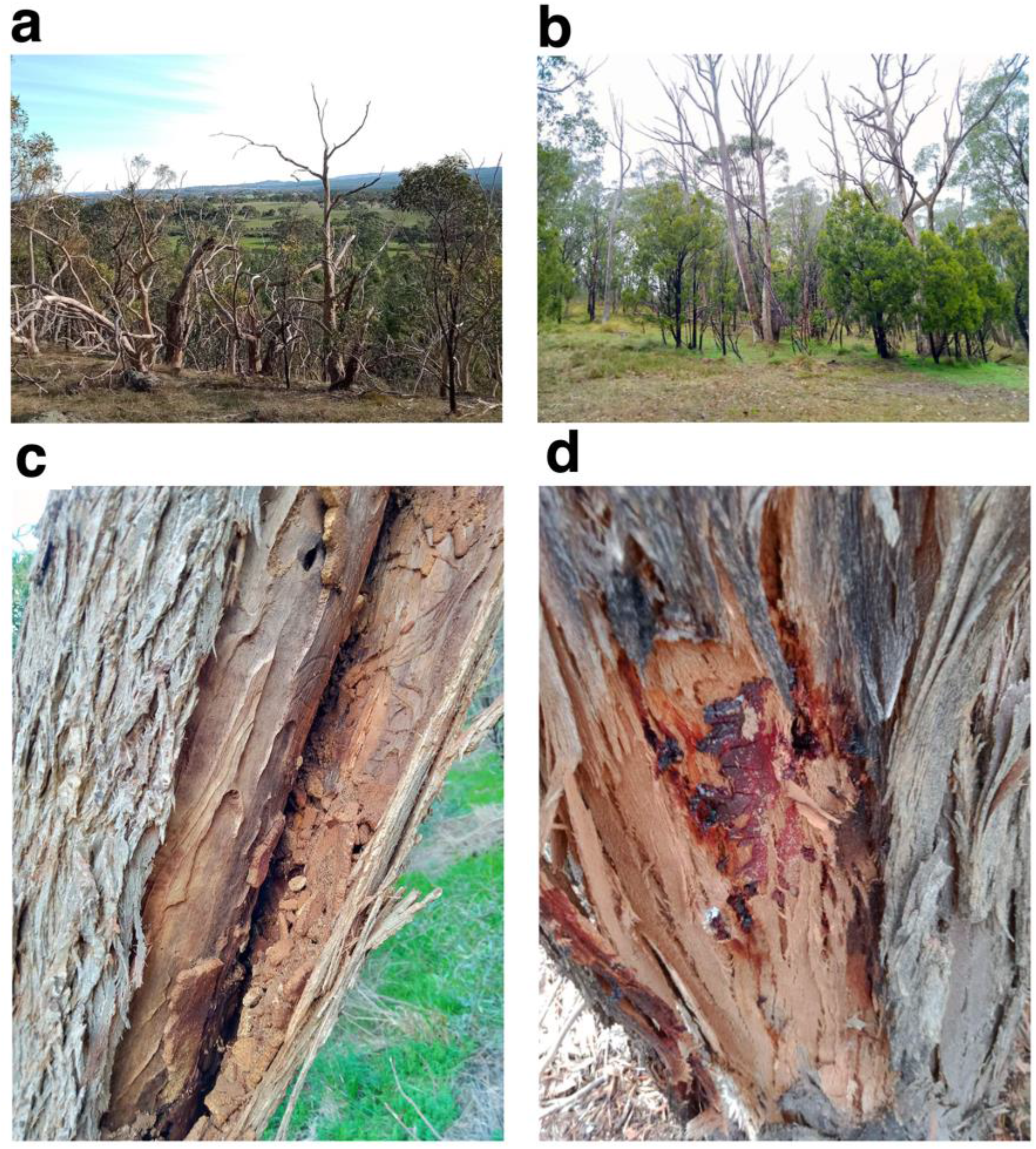
Selected photos from field assessment sites illustrating stringybark eucalypt dieback and its aetiology in the southern MLR. A) High mortality rate of *E. obliqua* on exposed ridge at Mount Pleasant summit, at the north-eastern edge of the species range; b) *E. obliqua* collapse site in Carey Gully (>1 m MAP) on a wet, south-facing slope and wet gully; c) signs of heavy borer infestation in dead tree such as dense larval galleries and frass under the bark; d) kino exudate in response to borer activity in living tree.

Dead trees were not obviously associated with particular habitats or environments, though such associations would need to be quantitatively tested with systematic sampling and statistical analysis. Patches of dead trees were seen in habitats including ridges, south facing slopes (Fig. 6), creek lines, deep soil in valley bottoms, shallow sand over rock with heathy understorey, and swamps. However, some of the most extensive mortality rates and heavily effected stands do include populations in the lower rainfall region of the species ranges, notably at locations such as Montacute, Mount Pleasant and Tanunda. Even so, collapsed stands were evident in higher rainfall areas of the Adelaide Hills.

PC infection was previously confirmed at one site that was inspected, ‘Hahndorf Soccer’. Here, mortality was approximately 80% (other tree species not affected). However, the trees had been ring-barked and killed by borers, not directly by PC. In general, mortality sites showed no obvious signs of PC. For example, leaf chlorosis was not observed, and susceptible species such as *Xanthorrhoea spp, Proteaceae spp*. etc were healthy whenever present in the understorey. However, this does not rule out the patchy presence and involvement of PC in tree mortality, especially as no laboratory tests for the presence of PC were conducted as part of these field investigations.

While many stands of stringybark have high density signs of leaf herbivore activity, other potential symptoms or signs such as leaf chlorosis, mistletoe infestation or other signs of ill health were generally not observed. Furthermore, while, as indicated by standing dead trunks without bark, there appears to have been some significant periods of tree mortality sometime within the past decade, several sites with relatively few dead trees had many living trees infested with borers (as detected through entrance holes with kino gum and signs of ring barking) that are likely to die in the near future.

### Analysis of existing tree monitoring data

Changes in tree health were assessed for stringybark sites with revisit data from BCM tree health module, in which the nearest 10 trees to the fixed centre-point are scored. Changes in canopy dieback status, proportion of dead stems and basal area were somewhat plot-specific. However, some patterns were evident. While in a few sites canopy dieback % decreased (indicating healthier trees), typically this metric increased between visits, indicated advancing canopy loss (Fig. 7). Indeed, canopy collapse >50% was recorded at three sites. In general, basal area increased modestly between surveys (most commonly up to 5%) while at a small number of sites, there was collapse with sharp declines in basal area due to tree mortality.

**Figure 7.**
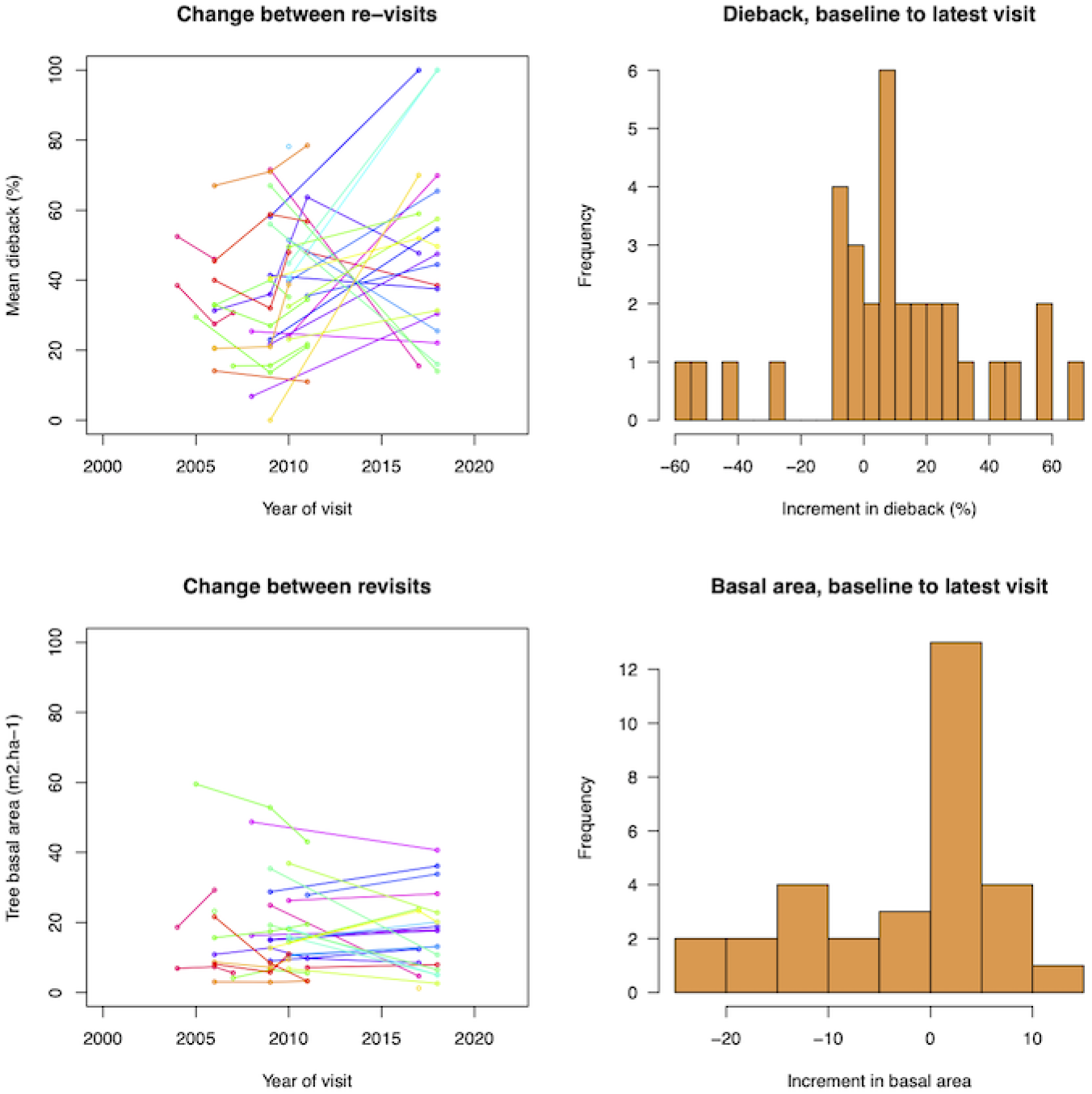
Trends and increments in canopy dieback and basal area based at BCM sites based on values averaged over the 10 trees surveyed at each site. Colours represent different stringybark sites with lines connecting baseline and subsequent visits. Basal area calculations are based only on the main trunk of multi-stemmed trees due to the BCM protocol and may not be an accurate representation of site-level increments.

### Conceptual model

We present two conceptual models of MLR stringybark forest dieback aetiology, one illustrating putative pathways for individual tree death and one illustrating pathways for widespread stand collapse across a landscape (Fig. 8). The pathways depicted are hypothesised based on literature and field observations and would need to be rigorously tested. We posit that stringybarks stressed by drought, disease (PC), leaf-feeding insects and fire are susceptible to borer infestations, which ultimately kill the trees. Processes leading to the death of individual trees emerge across the landscape, a scale at which contributing factors and patterns of tree mortality can be measured (e.g, topography and soils, climatic events, fire history, bird populations and plant functional traits).

**Figure 8.**
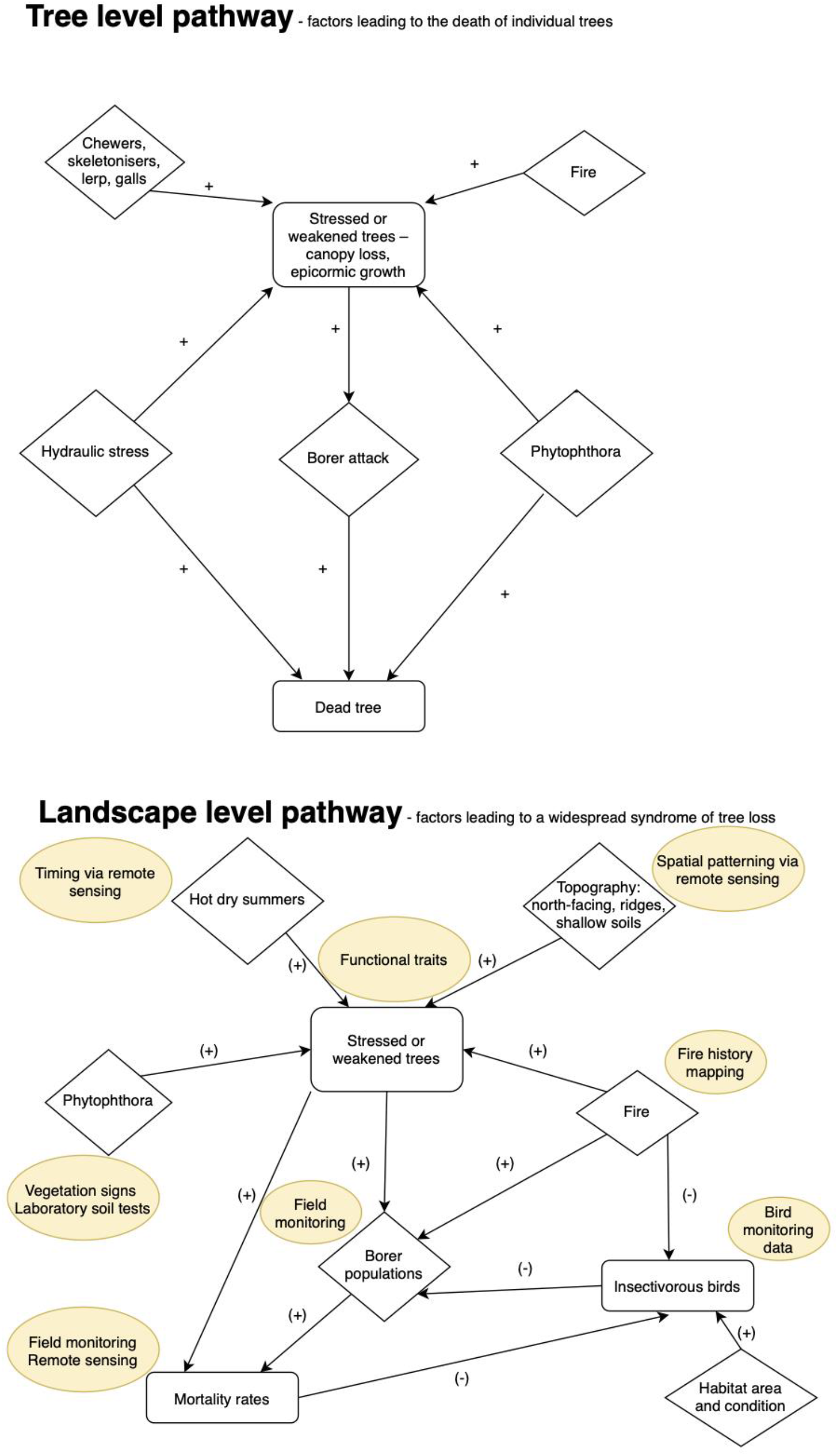
Schemas representing hypothetical pathways for the death of individual stringybark trees (*top*) as well as widespread syndromes of tree loss (*bottom*). Ovals in bottom panel highlight possible data/approaches to testing the importance of factors in landscape-level mortality rates.

## Discussion and Conclusion

### Concerning aetiology

Our desktop analysis of climatic niches shows that MLR populations of the two stringybark species, *E. baxteri* and *E. obliqua*, are spatially isolated from eastern Australian populations, and also experience higher summer aridity, presumably increasing vulnerability to heat and water stress, unless there has been local evolutionary adaptation to these conditions. Given that the climatic niche of stringybarks in the MLR is already outlying in terms of summer water deficit and aridity on long term averages, compared to eastern populations, incidences of drought years approximately 10 years apart with dry summers and high summer temperatures are likely to place these forests under particularly stressful conditions. Given climate change pressures (Pörtner et al. 2022), these conditions would logically be expected to be even more stressful in the drier sections of the species’ range, for example, Montacute– Cherryville and Barossa (e.g., Mount Pleasant Summit) regions for *E. obliqua*, and Kaiserstuhl for *E. baxteri*. Such climate ‘pinch-points’ (i.e., drought periods with summer heatwave conditions) suggest drought stress may cause or exacerbate the observed dieback in stringybark eucalypts. Indeed, *E. obliqua* has been reported to experience higher water stress during summer than co-occurring eucalypt species in the region (Sinclair 1980). If dieback were due to climate alone, future events should be relatively predictable with further data on hydraulic stress and the ability for trees to adjust physiologically and morphologically to repeated drought periods (Pritzkow et al. 2021).

In the case of *E. baxteri* and *E. obliqua*, however, more complex factors appear to be involved, potentially including the effects of past clearing, logging and fires, the resulting stand demographics, changes to avifauna, and the widespread, if patchy, presence of PC in wet forests of the region (Fig. S9), a disease to which both species are susceptible. Borers are implicated in tree mortality in these species based on field assessments at many sites and across the climatic range of the MLR distribution. Indeed, it is clear that, in most cases at least, it is borers rather than hydraulic failure per se that appear to be the ultimate cause of mortality. Larval feeding of vascular cambium tissue under the bark leads to ring-barking and death of trees (Seaton 2012). Previous stress caused by drought and other stressors weakens trees’ defences to such attacks (Schuldt et al. 2020), while the reallocation of nutrients from leaves to roots associated with crown senescence may increase the growth of phloem-feeding beetle larvae and contribute to outbreaks (White 2015). Seaton et al. (2015) found that longicorn beetles usually only attacked drought-stressed trees but that increasing drought frequency led to more frequent and severe outbreaks of these borers. Such outbreaks may result in the death of otherwise healthy trees. Longicorn (*Phoracantha* spp.) borers have caused stand collapse in Western Australia (Seaton et al. 2015) and cause problems in eucalypt plantations outside of Australia (Caldeira et al. 2002; Dhahri et al. 2016).

These observations tend to support a case that, for *E. baxteri* and *E. obliqua*, the causes of dieback consist of a complex landscape syndrome involving periods of drought stress combined with insect outbreaks and other local stressors that cause patchy stand collapse (see Fig. 8). The impact of this landscape syndrome may be moderated by disturbances and climate. For example, the frequency of disturbance such as fire may interact with disease and pest outbreaks to determine whether local populations can persist in the face of drought periods. (Yates et al. 2021). Furthermore, historically burnt trees that have resprouted from basal shoots to form multi-stemmed trees may be weakened and sometimes rot where the original trunk falls away. Therefore, there may be an imprint of past fires on current tree stress and dieback.

The aetiology of canopy loss and eventual mortality also determines the extent to which regeneration is possible and may vary across climatic gradients (Walters et al. 2005). While *E. baxteri* and *E. obliqua* are able to repair canopy from epicormic sprouts after drought or insect attack, and to regenerate from basal sprouts after fire, trees killed by heavy infestation of borers appear not to resprout at all. Borer attacks linked to drought stress may occur sometime after the event, meaning drought impacts have to be considered holistically as long term drivers of forest decline (Schuldt et al. 2020). Further refinement of our knowledge regarding the aetiology of stringybark dieback by species and region is needed before management implications can be fully understood.

Furthermore, the aetiology of, and susceptibility to, dieback may differ for Brown and Messmate Stringybark, and appear to be complex. Differences in vulnerability to dieback have also been reported for other eucalypt species (Ruthrof et al. 2015). In the Mount Lofty Ranges, Brown Stringybark is less affected by dieback (with the notable exception of Kaiserstuhl Conservation Park), though both species are susceptible to PC (Kueh et al. 2012). Furthermore, collapsed stands of Messmate Stringybark are consistently killed (at least in the end) by borers, whereas borers appear more incidental at the Brown Stringybark collapse site of Kaiserstuhl.

### Downstream consequences of stringybark dieback

Canopy collapse is a severe perturbation to the local environment, which causes cascading and rapid effects on ecosystem structure, composition and function (Anderegg et al. 2012; Pappas et al. 2022). Tree decline changes nutrient and water cycling and also the solar radiation and microclimatic environment at ground level (Wang et al. 2012). Canopy gaps favour light-loving species such as graminoids, and may benefit introduced weeds that thrive in more disturbed environments (Anderegg et al. 2012; Pappas et al. 2022). Reductions in the density of trees due to chronic disturbance can cause indirect loss of species richness due to environmental changes that resident species cannot tolerate (Jara-Guerrero et al. 2021).

Indeed, rapid changes in forest microenvironments can trigger thresholds for shifts between different vegetation types, while the loss of canopy as a buffer against extreme temperatures increases the exposure of resident species to climate change (Keppel et al. 2017). Angel & Bradley (2021) found direct links between declining rainfall and canopy loss in Wandoo eucalypt woodlands in Western Australia, and the abundance of canopy feeding birds was affected as a result. These downstream impacts to eucalypt forests suffering dieback and tree mortality are poorly documented for the target stringybark forests. A range of potential flow-on impacts (and benefits) could occur and are worthy of further investigation, for example, conversion of forest to heath, changes to microclimate, impacts on avifauna and shifts in the species composition of the understorey or overall diversity of the vegetation.

### Future research needed

Key future needs include understanding any differences in dieback aetiology among the species, documenting temporal and spatial patterns of dieback, quantifying dieback, and understanding the downstream effects of dieback. Closing these knowledge gaps will require ongoing and retrospective monitoring of existing populations. Currently, the existing network of Bushland Condition Monitoring sites form the best available baseline and could be used as the basis of further ground-based monitoring. While the 10 trees recorded in a given survey may not be sufficient to characterise stand demography in terms of distribution of age classes and accurate stand-level mortality rates, the method provides relatively rapid data, including distance between and bearing of trees, which can be used to calculate tree density (Elzinga et al. 1998). While the BCM method typically involves sampling just one point at a given monitoring site (where other vegetation condition parameters are recorded), it has the potential for generating meaningful indications of shifting parameters when multiple surveys in a region are pooled to create a larger sample. The dataset has the advantage of good baseline coverage of individual tree measures, which are sorely lacking in the study region as they are not included in most other survey methods used locally. Some methodological adjustments or clarifications may be required to maximise the utility of BCM for long term monitoring for this purpose, such as permanent tagging and measurement of all stems to monitor basal area.

A regional survey using remote sensing and ground-based techniques across a stratified network of sites, with nodes of more intense observation, would provide a much clearer picture of the extent and prevalence of mortality, canopy health status, and trajectory of basal area over time. More data on the landscape aetiology is required to understand the relationship of dieback events to habitat type, topography and differences in rainfall, to shape sensible site-specific management strategies. Field trials and vetting of management strategies techniques can then be contemplated, which we expect will encompass when and where to write-off forests, revegetation and regeneration strategies, and feasibility of borer treatment (Ministry of Forests, Lands and Natural Resource Operations 2012) and fire management.

### Conclusion

Canopy dieback and tree mortality are apparent in forests of the stringybark eucalypts *E. baxteri* and *E. obliqua* in the Mount Lofty Ranges of South Australia, with the latter more heavily affected. While the distribution of affected stands remains somewhat patchy across the landscape, there is potential for major ecosystem disturbance as a result of advancing canopy collapse and ensuing changes to forest microclimate and fauna. The dieback therefore threatens standing biodiversity and restoration programs in the region. Future research is needed to describe the full extent of the tree loss and to further investigate contributing factors, including pathogens, water stress associated with drought periods and heat waves, and their interaction with soil and topography, insect attacks, altered fire regimes, altered ground water table levels, past logging, changes to faunal assemblages, and complex syndromes involving multiple interacting factors. Early intervention strategies may focus on regeneration as well as fire and pest management.

## Acknowledgements

We thank Trees For Life Inc. and the Stringybark Dieback Steering Group (Adelaide Hills Council, Mount Barker Council, Northern & Yorke Landscape SA Board, Hills & Fleurieu Landscape SA Board, and the Nature Conservation Society of South Australia) who provided funding, data and advice for this work. We also thank participating stakeholders, students and experts, including Rachael Nolan, and the South Australian Department for Environment and Water.

## Additional figures

Figure S9 – *Phytophthora cinnamomi* (PC) map.

Figure S10 – SPEI index time series.

**Figure S9.**
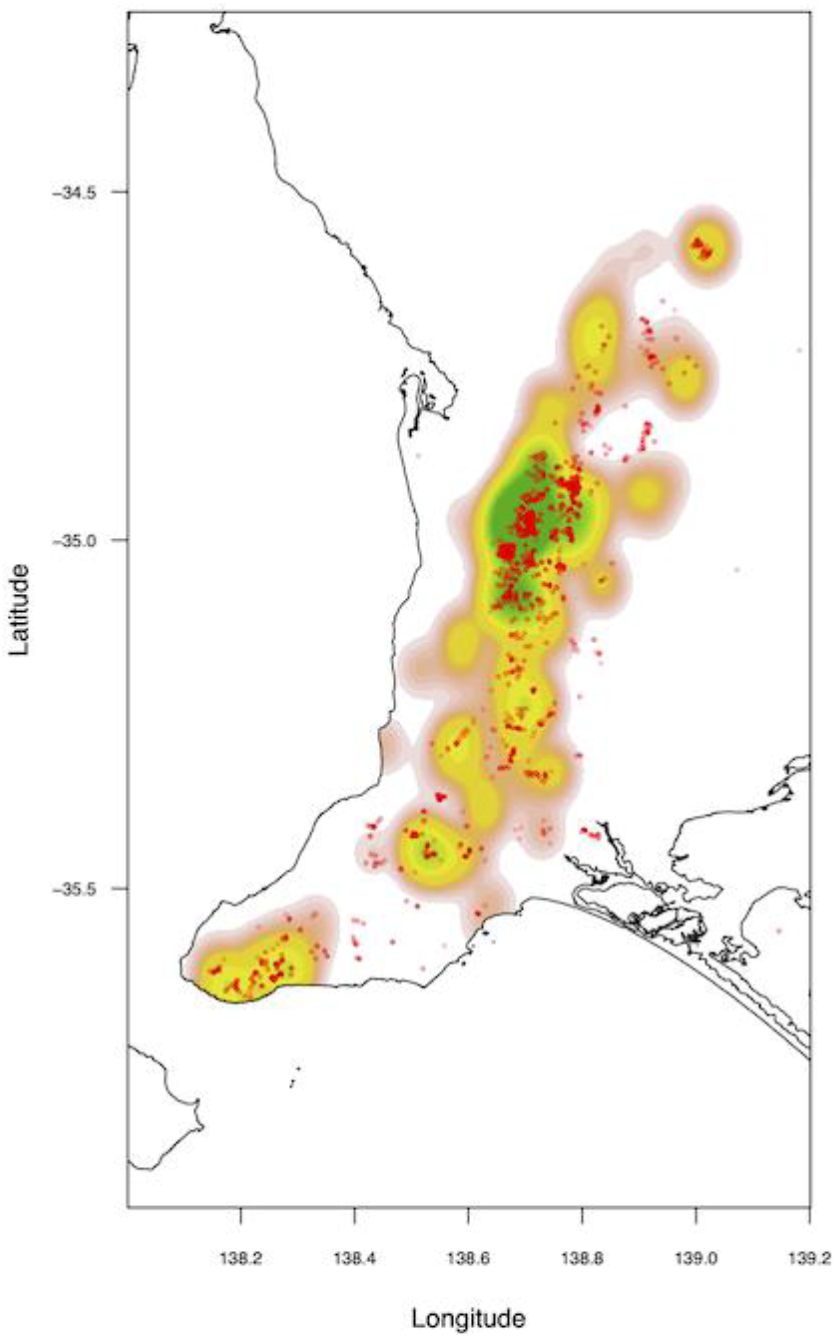
Spatial kernel density of *Phytophthora cinnamomi* (PC) sites in the study region, showing high correspondence to densities of stringybark records (overlayed points).

**Figure S10.**
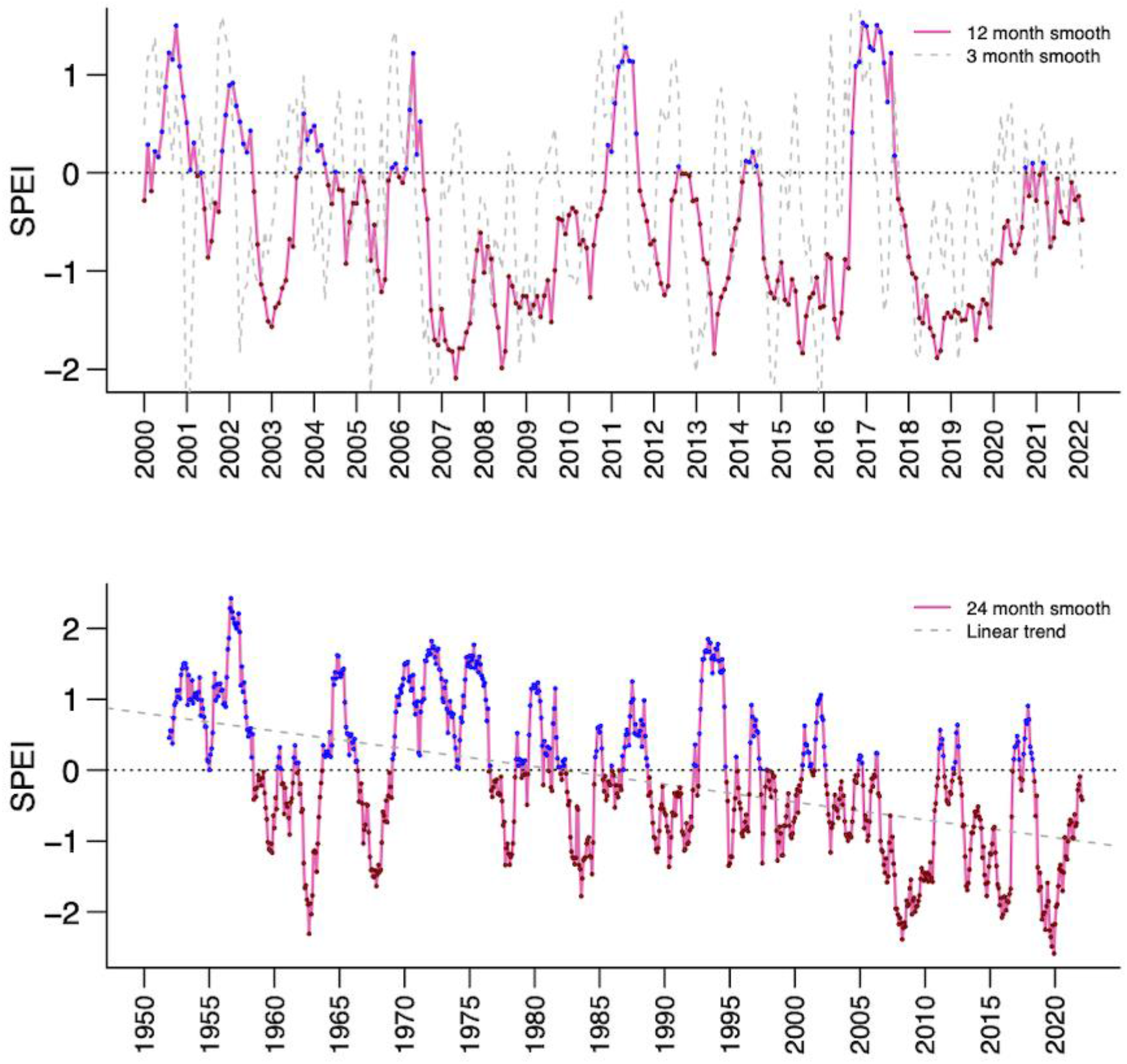
Standardised Precipitation-Evapotranspiration Index anomaly (source global SPEI database) for the study region. Above: quarterly and yearly averaged data since 2000, showing drought periods such as 2006– 2009, 2015 and 2018–2019. Below: 2-yearly averaged data since 1950, showing significant droughts (2006– 2009 and 2019) and an overall trend towards increasing frequency of drought periods relative to the long-term mean (especially notable 2000–).

## Notes

### Competing Interest Statement

The authors have declared no competing interest.

